# Neuroimaging genomics as a window into the evolution of human sulcal organization

**DOI:** 10.1101/2023.10.23.563571

**Authors:** Ole Goltermann, Gökberk Alagöz, Barbara Molz, Simon E. Fisher

**Author notes:** These authors contributed equally to this work (Ole Goltermann, Gökberk Alagöz).

## Abstract

Primate brain evolution has involved prominent expansions of the cerebral cortex, with largest effects observed in the human lineage. Such expansions were accompanied by fine-grained anatomical alterations, including increased cortical folding. However, the molecular bases of evolutionary alterations in human sulcal organisation are not yet well understood. Here, we integrated data from recently completed large-scale neuroimaging genetic analyses with annotations of the human genome relevant to various periods and events in our evolutionary history. These analyses identified single-nucleotide polymorphism (SNP) heritability enrichments in foetal brain human-gained enhancer elements for a number of sulcal structures, including the central sulcus, which is implicated in human hand dexterity. We zeroed-in on a genomic region which harbours DNA variants associated with left central sulcus shape, a human-gained enhancer element, and genetic loci involved in neurogenesis including *ZIC4*, to illustrate the value of this approach for probing the complex factors contributing to human sulcal evolution.

---

The evolution of mammals led to considerable interspecific diversity in neuroanatomy, including substantial cortical expansions in the primate order (Herculano-Houzel 2011). These expansions were accompanied by an array of changes at different anatomical levels, such as the emergence of primate-specific cell types (Schmitz et al. 2022), overall increases in cortical surface area (Van Essen et al. 2018), and reorganisations of gyral and sulcal morphology (Amiez et al.). As humans have the highest gyrification index among primates (Mota and Herculano-Houzel 2015), the gyral and sulcal organisation of the human cortex stands out as a trait of particular evolutionary interest, with potential relevance for aspects of cognition and behaviour (Amiez et al. 2018; Willbrand et al. 2022). Although the anatomical basis of sulcal morphology in primates is rather well-studied (Ronan and Fletcher 2015), we are only just beginning to gain insights into genetic pathways that contribute to overall degree of gyrification, and regional distribution and morphology of individual sulci. Moreover, the ways in which such genetic pathways changed in hominin evolution are poorly understood.

While comparative approaches have revealed critical differences in gyral and sulcal organisation across primates (Mota and Herculano-Houzel 2015; Amiez et al. 2019; Amiez et al. 2021), findings are mainly limited to gross anatomical features, and do not provide insights into their biological bases. On the other hand, much of our understanding of molecular genetic contributions to primate brain evolution comes from *in vitro* neuronal and organoid models, and *in silico* comparative genomic and transcriptomic studies, which have identified primate-specific brain cell-types and human-specific molecular changes (Lodewijk et al. 2020; Schmitz et al. 2022). Yet, such findings often require validation in primary tissue, which is strictly limited by the scarcity of high-quality non-human primate brain samples. Thus, there remains a large gap between the comparative neuroimaging and experimental approaches, hindering a comprehensive understanding of human and non-human primate brain evolution.

Advances in large-scale neuroimaging genomics investigations in present-day humans have lately proven useful to bridge this gap, as they combine some aspects of neuroanatomical and molecular approaches. Datasets encompassing both brain magnetic resonance imaging (MRI) and single-nucleotide polymorphism (SNP) information from tens of thousands of people, such as the UK Biobank (Elliott et al. 2018), and the Enhancing Neuro Imaging Genetics through Meta-Analysis (ENIGMA) Consortium (Grasby et al. 2020), have allowed the first well-powered genome-wide association studies (GWAS) of interindividual differences in a variety of neuroanatomical traits including cortical surface area, white-matter connectivity, and sulcal morphology (Shadrin et al. 2021; Zhao et al. 2021; Brouwer et al. 2022). In parallel, the fields of ancient DNA and comparative primate genomics have been generating considerable amounts of genomic and functional data, which highlight regions of the genome that may have been important in human evolution. Availability of multiple high-quality Neandertal genomes (Prüfer et al. 2014; Prüfer et al. 2017; Mafessoni et al. 2020) led to the identification of Neandertal introgressed alleles in present-day humans as a result of the admixture events from ∼50-60,000 years ago (Vernot and Akey 2014). This enabled genomic annotations not only of Neandertal introgressed fragments, but also of regions that are significantly depleted of Neandertal ancestry, *archaic deserts*, in present-day humans (Vernot et al. 2016). On the other hand, comparing histone modification profiles of foetal human, macaque and mouse brain tissues revealed regions of the genome that are active solely in the foetal human brain, referred to as *human-gained enhancers* (HGEs), with potential relevance within the last ∼30 million years of evolution along the lineage that led to *Homo sapiens* (Reilly et al. 2015).

Initial work by Lemaitre et al. (2023) investigated the evolution of sulcal opening and depth using summary statistics from 18,000 individuals of European ancestry from UK Biobank, in combination with evolutionary annotations of the human genome compiled by Tilot et al. (2021). While they found significant SNP-heritability enrichment for foetal brain HGEs in left and right calloso-marginal posterior fissures and the right central sulcus, their approach also showed a number of significant partitioned SNP-heritability estimates that had a negative sign, a pattern of findings which is biologically implausible. They highlighted that these might reflect limitations of the study related to i) low SNP proportions covered by evolutionary annotations relative to the reference panel, and ii) the sample size of the early UK Biobank neuroimaging data release (Bycroft et al. 2018), which was relatively modest for contemporary GWAS efforts. In this study, we adopted an improved analytic pipeline to address these limitations by using three newly curated evolutionary annotations with sufficient SNP-coverage (Alagöz et al. 2022), two additional sulcal morphology measures that were not investigated by Lemaitre et al. (2023) (sulcal length and surface area), and an even larger sample size.

Here, we made use of results from the latest genome and exome wide association study on regional sulcal morphology in around 26,530 individuals in UK Biobank, which incorporated four regional shape parameters (length, mean depth, surface area and width) from each sulcus, comprising in total 450 sulcal parameters across 58 sulci (Sun et al. 2022). We integrated the summary statistics of this study with an enhanced set of evolutionary annotations, and split our analytic pipeline into two parallel streams: 1) a hypothesis-based targeted study of four sulci with high evolutionary relevance based on prior literature, and 2) a hypothesis-generating exploratory study of 45 sulci, with both streamlines focusing on regional hemispheric sulcal anatomy measures. For the targeted study, we hypothesised that the common genetic variation underlying present-day anatomic variation in these sulci would be enriched in foetal brain HGEs, as the sulci are phenotypically divergent between humans and Old World monkeys (OWMs). In addition, given the variable organisation of such sulci across apes, we might expect enrichment or depletion signals in evolutionary annotations that tag more recent timescales, such as those related to Neandertal ancestry.

The four sulci of interest in our targeted approach (the central sulcus, the paracentral sulcus, the parieto-occipital sulcus, and the superior temporal sulcus), are important and relevant for human brain evolution for a number of reasons. The central sulcus was shown to exhibit changes in surface area, and in its folding pattern, particularly during the evolution of OWMs by a comparative study using magnetic resonance imaging of OWM, apes and humans (Patel et al. 2019). The paracentral sulcus (Amiez et al. 2019), the parieto-occipital sulcus (Hopkins et al. 2014), and the superior temporal sulcus (Bruner 2018) show the most marked morphological changes when comparing OWMs to apes and humans. For the targeted approach, we aimed to exploit all four available sulcal shape descriptors, and identified significant SNP-heritability enrichment signals in human-gained enhancers for the width of both the right and left central sulcus, extending prior right-hemispheric findings (Sun et al. 2022). The exploratory stream involved analyses of 45 sulci, and revealed that common genetic variants associated with the right hemisphere olfactory sulcus depth are significantly enriched in HGE elements. We demonstrate that combining neuroimaging genetics with heritability-based post-GWAS evolutionary analyses can shed light on aspects of human brain evolution by identifying links between brain structures and evolutionary annotations of the human genome, and highlighting genes of particular interest for future experimental follow-up.

## Materials and Methods

### GWAS Summary Statistics

Sulcal morphology GWASs were performed by Sun et al. (2022) using a recent release of the UK Biobank brain imaging and genotype data set (N=26,530). Specifically, 62 sulcal folding traits per brain hemisphere and four sulcal shape descriptors were examined, resulting in 450 sulcal parameter phenotypes after excluding shape descriptors with >75% missingness, including 58 sulci and four sulcal shape parameters. The brain imaging dataset contains reliable quantifications of four sulcal shape descriptors: sulcal depth, length, width and surface area, each representing a different aspect of sulcal morphology. Imaging phenotypes were inverse-rank normalised in order to approximate a standard normal distribution, and minimise outlier effects. The total sample of 40,169 individuals was split into discovery (N_Discovery_=26,530) and replication (N_Replication_=13,639) cohorts, with the discovery cohort including data from only European ancestry individuals and the replication cohort including admixed ancestries, which would likely confound any heritability-based evolutionary analyses (Browning and Browning 2011). Sample sizes differed depending on the specific sulcal parameters, and are listed in Tables S2 and S3. The online browser for GWAS summary statistics is publicly available at https://enigma-brain.org/sulci-browser (see Sun et al. (2022) for details).

### SNP-heritability thresholding

We performed univariate LDSC (Bulik-Sullivan et al. 2015) to estimate total SNP-heritability of 450 sulcal parameters obtained from Sun et al. (2022) GWAS summary statistics to optimise our partitioned heritability analysis. We then filtered sulcal parameters based on these SNP-heritability estimates in using R (v4.0.3), prior to performing LDSC partitioned heritability analysis. Our filtering process consisted of two stages: 1) exclusion of traits with SNP-heritability <10%, and 2) exclusion of traits with non-significant (α_Bonferroni_=0.05/163) SNP-heritability estimates. We performed the significance test by generating the cumulative probability distribution of the total SNP-heritability estimates of 163 pre-selected sulcal parameters, using the *p.norm* function of R-base (v4.0.3), yielding 153 sulcal parameters with significant SNP-heritability estimates for the subsequent partitioned heritability analysis.

### Estimating Number of Independent Neuroimaging Traits

For the targeted and exploratory analysis streams, we reduced the multiple-testing burden by considering the genetic correlations across investigated traits. We used spectral decomposition of matrices (SpD) implemented in the R package PhenoSpD (Zheng et al. 2018) to estimate the effective number of independent variables (VeffLi). This resulted in 9.48 and 73.59 independent traits for the targeted and exploratory analysis subsets, respectively.

### Partitioned Heritability Analysis

Contributions of evolutionary annotations to the total SNP-heritability of each sulcal morphology measure were computed using the LDSC partitioned heritability tool (Finucane et al. 2015), following the guideline and tutorials in the LDSC Github Wiki website (https://github.com/bulik/ldsc/wiki/PartitionedHeritability). We exploited three evolutionary annotations prepared by Alagöz et al. (2022), with increased SNP-coverage ratios covering at least 1% of the SNPs in the 1000 Genomes reference panel (Auton et al. 2015) (HapMap3 SNPs, the MHC region SNPs were excluded): Foetal brain HGEs (Reilly et al. 2015), Neandertal introgressed alleles (Vernot and Akey 2014), and archaic deserts (Vernot et al. 2016). All annotations were controlled for the baselineLD v2.2 model provided by the Alkes Price’s group, whereas the foetal brain HGE annotation was additionally controlled for the active regulatory elements from the Roadmap Epigenomics Consortium database (Ernst and Kellis 2012). The foetal brain HGEs annotation covers the enhancer elements that are active in human foetal cortical brain tissue at 7^th^, 8.5^th^ and 12^th^ post-conception weeks, while inactive in macaque and mouse brains during the corresponding developmental stages. The Neandertal introgression annotation is a list of SNPs including the genomic coordinates of the introgressed SNPs, and their “LD-friends” (SNPs that are in perfect LD (*r^2^*=1) with the introgressed allele). The archaic deserts annotation encompasses large stretches of the genome that are significantly depleted for Neandertal introgressed alleles. These deserts were identified by Vernot et al. (2016) by using a sliding window approach, which detects regions that are 10Mb or larger and significantly depleted of Neandertal DNA in various populations including Europeans, Asians, and Melanesians (average introgression percent per region<10^-3.5^, see Vernot et al. (2016) for details).

### Functional and Evolutionary Annotation of GWAS Loci

For left hemisphere central sulcus width, we performed a qualitative analysis to identify overlaps between foetal brain HGE elements and genome-wide significant (P<5×10^-8^) loci (*r^2^*>0.6), which we identified using the clump function of PLINK (Purcell et al. 2007). After detecting the genome-wide significant SNPs associated with width of the left central sulcus that fall into the foetal brain HGE annotation, we focused on an example region harbouring a GWAS hit, an HGE element and *ZIC4*/*ZIC1* genes. We used the SNP2GENE function of Functional Mapping and Annotation of Genome-Wide Association Studies (FUMA, version 1.3.6a, Watanabe et al. 2017) to assess the chromatin interaction profile of the *ZIC4* locus with data from foetal and adult brain samples (Schmitt et al. 2016; Giusti-Rodríguez et al. 2019). We investigated the cis-eQTL properties of the GWAS hit lead SNP using the MetaBrain brain tissue gene expression database (Marzke 1997). For the comparative genomic analysis, we derived the Neandertal and Denisovan allelic states for our SNP of interest using the publicly available ancient genotype dataset by the Max Planck Institute for Evolutionary Anthropology (cdna.eva.mpg.de/neandertal/), and used the Ensembl phylogenetic context server (Howe et al. 2021) to fetch the allelic states of non-human primates.

## Results

### SNP-heritability Estimation and Establishment of The Targeted and Exploratory Analytic Streams

We used publicly available sulcal morphology GWAS summary statistics (UK Biobank, N=26,530) of 450 sulcal parameters (see Sun et al. (2022) for the details). These GWASs were performed on 58 sulci across brain hemispheres, and four sulcal shape descriptors (mean depth, surface area, width, length). We estimated the total SNP-heritability of each sulcal parameter using univariate linkage-disequilibrium score regression (LDSC, Bulik-Sullivan et al. 2015), and showed that the SNP-heritability estimates varied across sulci, and sulcal shape descriptors ranging between 0.07-31%, with a median SNP-heritability of 7%(SE=0.03) (Fig. 1A, Table S1).

**Fig. 1:**
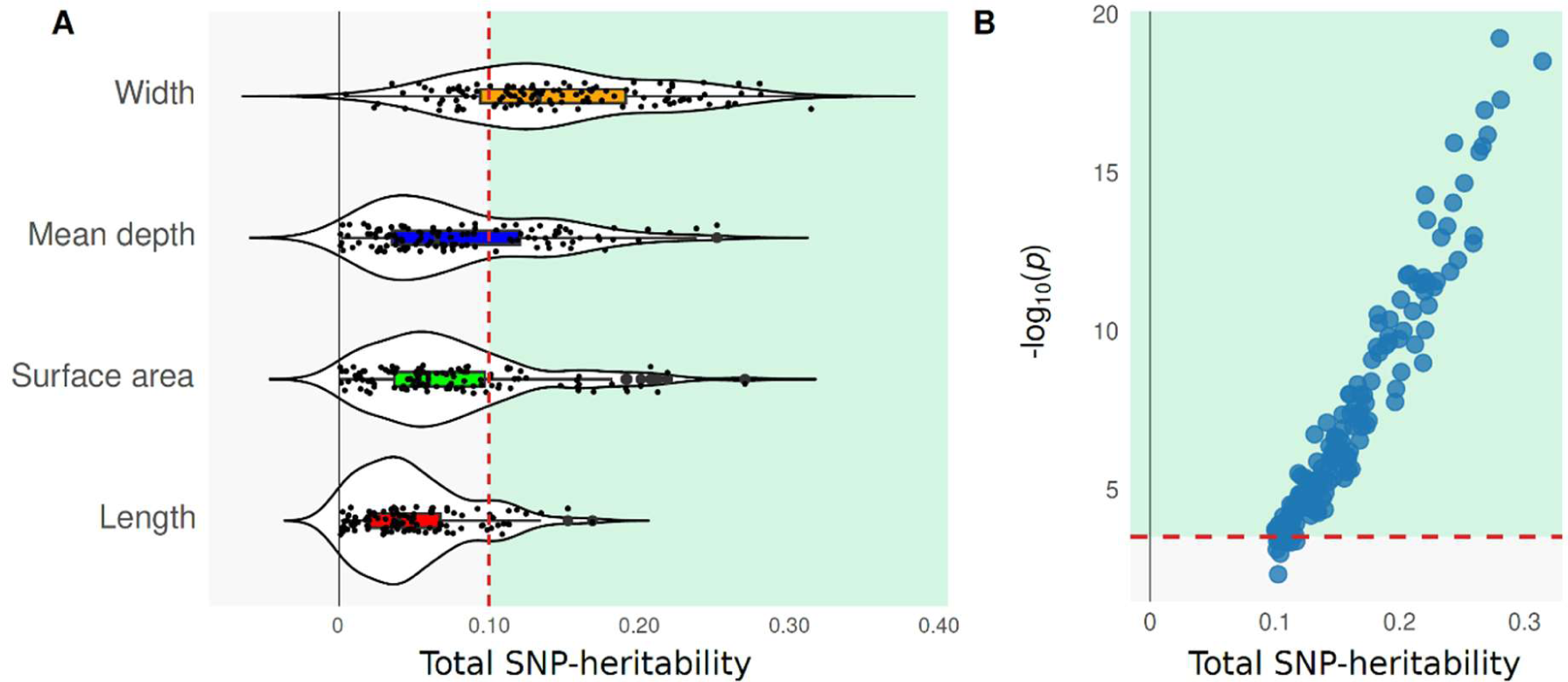
Univariate SNP-heritability estimates and heritability-based filtering of sulcal parameters. (A) Total SNP-heritabilities of the 450 sulcal parameters estimated using the GWAS summary statistics from Sun et al. (27). The mean total SNP-heritability estimates varied across sulcal shape descriptors. Traits with >10% heritability were kept for the second step of filtering. Red dashed line indicates the 10% cut-off. Traits that are kept for subsequent filtering are highlighted in green, filtered ones are depicted in grey. (B) - log_10_(*p*-values) reflecting SNP-heritability significance of each sulcal parameter. Red dashed line indicates the significance threshold (-log_10_(0.05/163)). Traits that are kept for partitioned heritability analysis are highlighted in green, filtered ones are depicted in grey.

Given the importance of trait heritability for obtaining statistically reliable partitioned heritability estimates (Finucane et al. 2015), we optimised our analytic pipeline for the subsequent partitioned heritability analysis (i.e. to minimise negative partitioned heritability estimates, reduce the standard errors, and minimise the multiple-testing burden). For this purpose, we applied a two-step heritability-based filtering, before performing any enrichment tests. We first filtered out traits with a SNP-heritability estimate of less than 10% (Tashman et al. 2021), and obtained 163 sulcal parameters (52 sulci, four sulcal shape descriptors) (Fig. 1A). Secondly, we excluded any trait for which the SNP-heritability estimate did not meet significance (α_Bonferroni_=0.05/163, significance test against zero), yielding 153 sulcal parameters (49 sulci, four sulcal shape descriptors) (Fig. 1B). These traits were used to curate two subsets for our targeted and exploratory analysis streams. For the targeted analysis, we included the central sulcus, the paracentral sulcus, the parieto-occipital sulcus, and the superior temporal sulcus, and three sulcal shape descriptors. These criteria yielded, across the two hemispheres, a total of 14 sulcal parameters with sufficient SNP-heritability (range: 12-31%, four sulci, three remaining sulcal shape descriptors) (Table S2). Importantly, none of the sulcal length parameters survived our heritability thresholding for the targeted trait selection. For the exploratory stream, we did not pre-select any sulci, and included all four shape descriptors again. Consequently, 139 sulcal parameters with sufficient SNP-heritability estimates were identified for the exploratory analysis stream (range: 10-28%, 45 sulci, four sulcal shape descriptors) (Table S3).

As shown by Sun et al. (2022), some of the sulcal parameters are phenotypically and genetically correlated across and within brain hemispheres. Hence, we estimated the effective number of independent traits within targeted and exploratory stream subsets using PhenoSpD (v1.0.0, Zheng et al. 2018), and two genetic correlation matrices of 14 and 139 sulcal traits, which identified nine and 74 independent variables, respectively. The number of independent variables was subsequently used for multiple testing correction.

### Central Sulcus Width-associated Genetic Variants are Enriched in Foetal Brain HGEs

In the targeted study, we performed LDSC-based partitioned heritability (Finucane et al. 2015) analysis of the 14 pre-selected sulcal parameter GWAS summary statistics from UK Biobank (N=26,530) (Sun et al. 2022) that showed sufficient heritability estimates (according to filtering criteria described above), with a median of 16% (Table S2). For each of these sulcal parameters, we tested SNP-heritability enrichment/depletion in three evolutionary annotations covering different evolutionary timescales: foetal brain HGEs, Neandertal introgressed alleles, and archaic deserts. After false discovery rate (FDR) correction for nine independent sulcal parameters, we identified significant SNP-heritability enrichments in foetal brain HGEs for the width of left and right central sulci (left: Enrichment(SE)=7.75(2.52), P_FDR_=0.04, right: Enrichment(SE)=7.57(2.44), P_FDR_=0.04) (Fig. 2, Table S4), as well as right parieto-occipital sulcus surface area (Enrichment(SE)=13.03(3.86), P_FDR_=0.01), and right superior temporal sulcus width (Enrichment(SE)=5.66(1.94), P_FDR_=0.04) (Fig. 2, Table S4). No other significant positive enrichments or depletions were found in Neandertal introgressed alleles and archaic deserts annotations for the remaining sulcal parameters in our targeted analysis (Table S5-6).

**Fig. 2:**
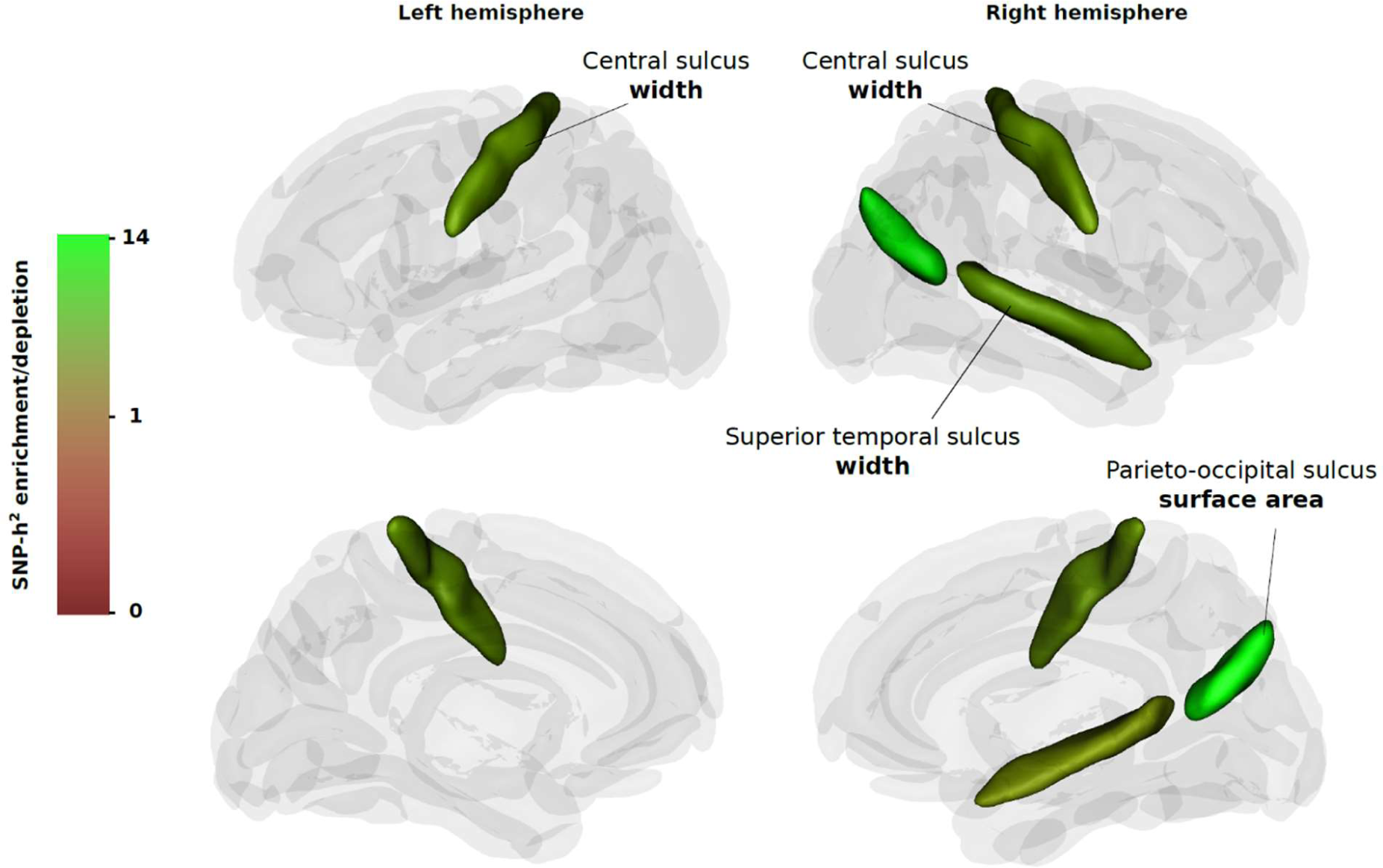
Three-dimensional visualisation of brain sulcal structure and SNP-heritability enrichment/depletion levels in foetal brain HGEs (covering active enhancer elements at 7^th^, 8.5^th^, and 12^th^ post-conception weeks) for the targeted sulci in each hemisphere. Non-significant results for targeted sulci are shown in dark grey, other sulci in light grey.

### Right Hemisphere Olfactory Sulcus Width SNP-heritability is Enriched in Foetal Brain HGEs

For the exploratory stream we widened the scope of our analysis to examine sulcal morphology across the whole brain. We tested the 139 sulcal trait GWAS summary statistics from UK Biobank (N=26,530) (27) with a median SNP-heritability estimate of 15%(SE=0.03) for SNP-heritability enrichment and depletion in the same three evolutionary annotations as the targeted approach. After FDR correction for 74 independent sulcal parameters, we identified a significant heritability enrichment in foetal brain HGEs for the mean depth of the right olfactory sulcus (Enrichment(SE)=11.26(2.91), P_FDR_=0.02) (Fig. 3, Table S7). We also identified significant SNP-heritability depletions in the archaic deserts for the width measures of five sulcal parameters: The left internal frontal sulcus (Enrichment(SE)=0.11(0.25), P_FDR_=0.03), the left inferior frontal sulcus (Enrichment(SE)=0.07(0.32), P_FDR_=0.04), the left posterior inferior temporal sulcus (Enrichment(SE)=0.13(0.30), P_FDR_=0.04), as well as the right insula (Enrichment(SE)=0.28(0.24), P_FDR_=0.04), and the right posterior terminal ascending branch of the superior temporal sulcus (Enrichment(SE)=0.14(0.29), P_FDR_=0.04) (Fig. 3, Table S8). We did not detect any significant positive enrichment or depletion signals for the Neandertal introgressed alleles annotation (Table S9).

**Fig. 3:**
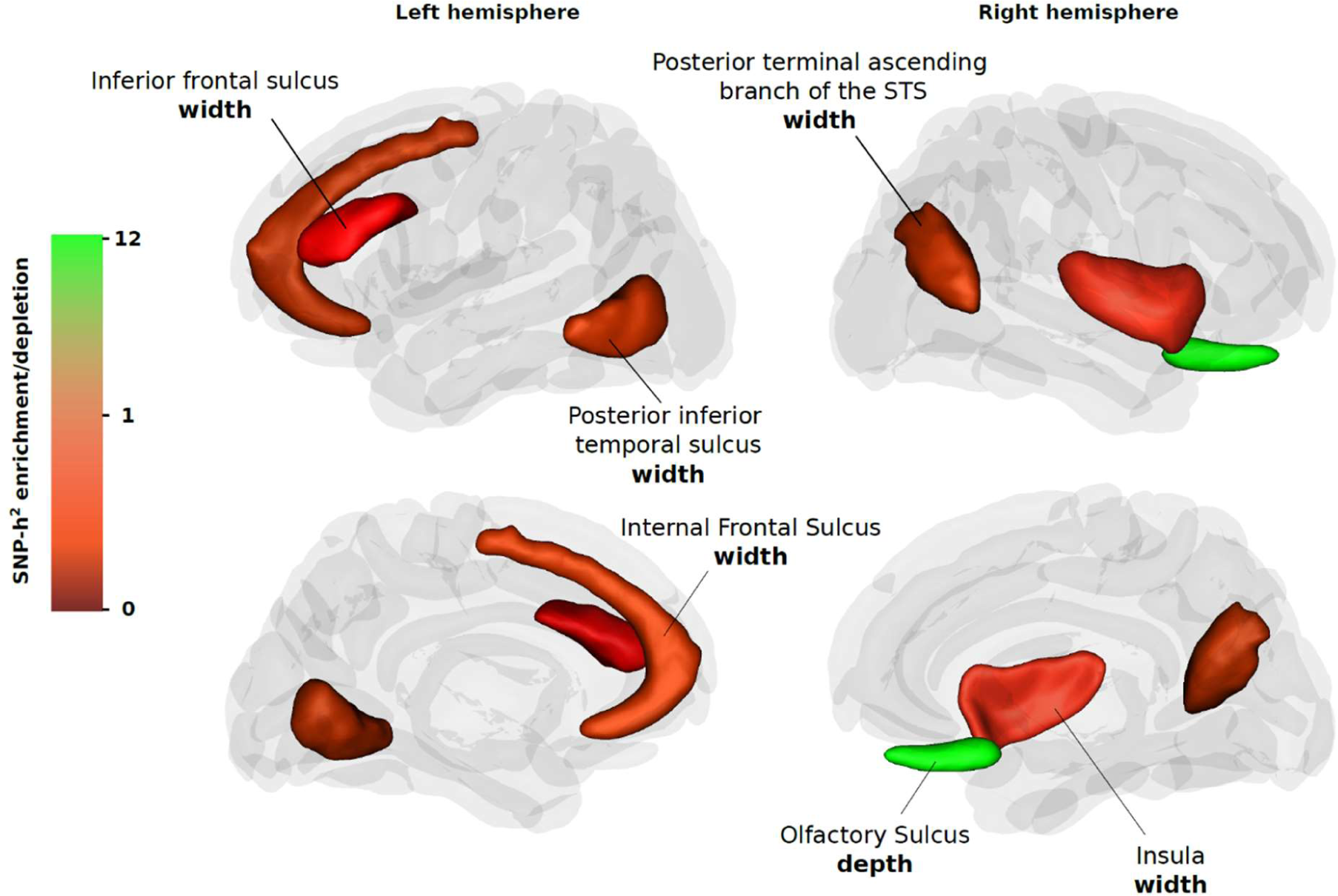
Three-dimensional visualisation of brain sulcal structure and SNP-heritability enrichment/depletion levels in foetal brain HGEs and archaic deserts for the sulci analysed in the exploratory analysis stream. Non-significant results for targeted sulci are shown in dark grey, other sulci in light grey.

### Left Central Sulcus Width-associated Variants Overlap with an HGE Co-localized with *ZIC4* and *ZIC1* Genes

Finally, we aimed to pin down putative loci and genes implicated in the evolution of human sulcal organisation by evaluating the links among sulcal morphology-associated common genetic variation, HGEs, and chromatin interaction patterns in foetal human brain tissue. Given the importance of the central sulcus in manual motor skills and dexterity, which are critical human adaptations (Marzke 1997; Tocheri et al. 2008), and the fact that it yielded two robust significant SNP-heritability enrichment signals (Table S4), we followed up with evolutionary and functional annotations of the common genetic variants associated with this sulcus. Out of 11 independent genome-wide significant (P<5×10^-8^) loci (blocks of LD covering lead SNPs and adjacent SNPs in linkage disequilibrium, *r^2^*>0.6) associated with left central sulcus width, we detected five LD-blocks that overlap with foetal brain HGE elements (Table S10). Intriguingly, one of these five loci is tagged by the independent genome-wide significant SNP rs884370 (P=1.39×10^-8^, Beta(SE)=-0.05(0.009)), which is located in a foetal brain HGE element adjacent to the *ZIC4* gene (chr3:147,101,875-147,103,850) (Fig. 4A). We further annotated the locus using the chromatin interaction data from foetal brain tissue to test the regulatory relevance of this enhancer element, and the SNP rs884370. FUMA functional annotation (Watanabe et al. 2017) revealed that the HGE element interacts with the promoter regions of *ZIC4* and *ZIC1* (Fig. 4A).

**Fig. 4:**
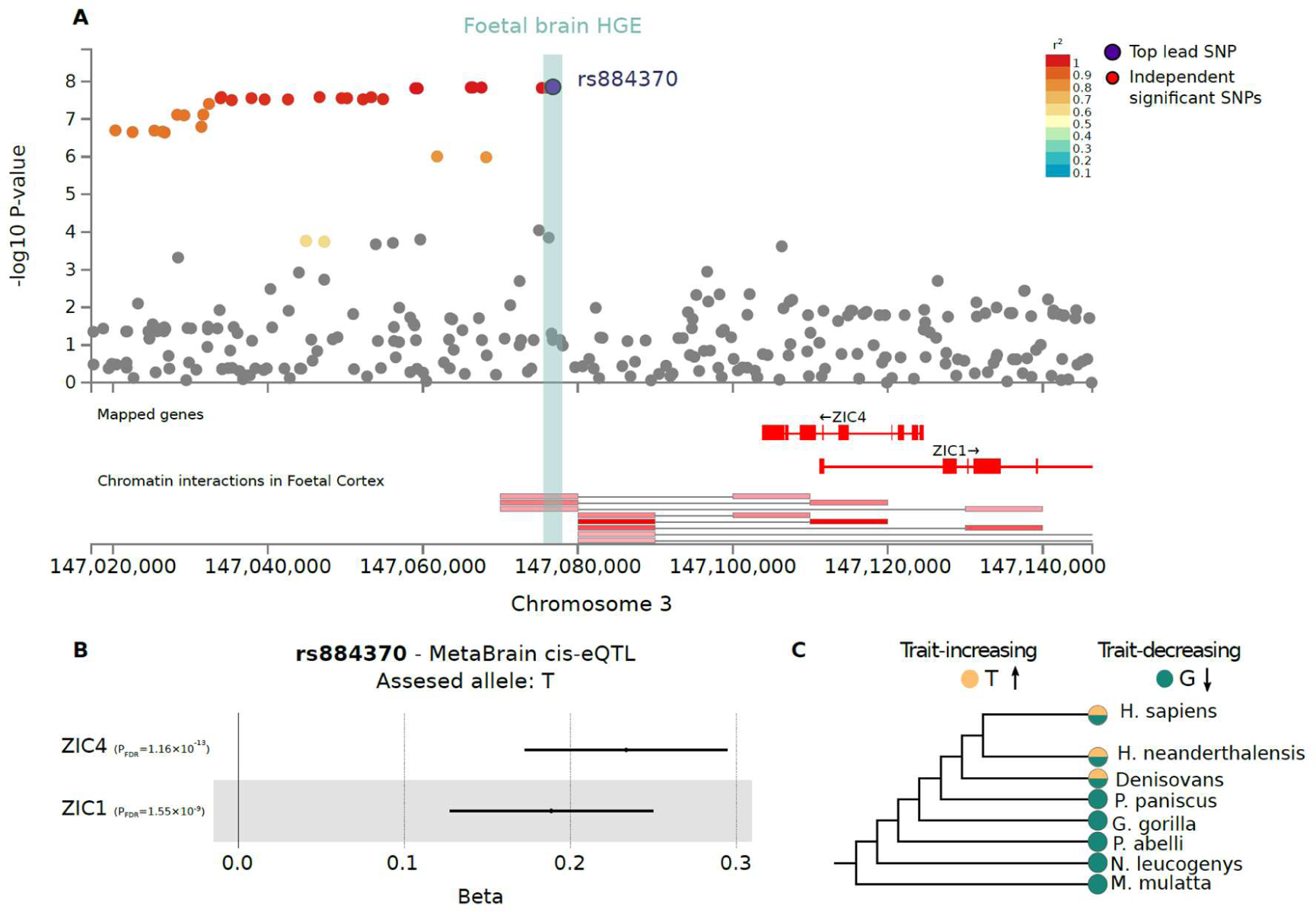
Functional-evolutionary annotation of the locus around rs884370, which is associated with the width of the left-hemisphere central sulcus. (A) The LocusZoom plot obtained by FUMA shows SNPs with genome-wide significant association (P<5×10^-8^). The foetal brain HGE element which overlaps with rs884370 is shown as a light green bar. Colours represent linkage disequilibrium with the lead SNP rs884370. The chromatin interaction profile of the locus demonstrates the overlap between the HGE element and the interaction region. hg19 coordinates are shown. (B) cis-eQTL plot of rs884370, showing the impact of this SNP on *ZIC4* and *ZIC1* expression in adult brain tissue for the T allele. Error bars represent the standard error. (C) Derived and ancestral allelic states for rs884370 are shown for *H. sapiens*, Neandertals, Denisovans and five non-human primates. Colour coding indicates the derived and ancestral states.

We further scrutinised the regulatory and evolutionary relevance of the SNP rs884370 by leveraging the MetaBrain expression quantitative trait loci (eQTL) dataset (de Klein et al. 2023), a publicly available ancient DNA genotype dataset, and non-human primate genome assemblies. Our analyses revealed that rs884370 is annotated as a cis-eQTL of *ZIC4* (P=5.81×10^-14^, Beta(SE)=0.23(0.03)) and *ZIC1* (P=1.55×10^-14^, Beta(SE)=0.23(0.03)) in adult brain tissue (Fig. 4B, Table S11). The assessed (alternative) allele of rs884370 (T) is associated with increased expression of both *ZIC4* and *ZIC1* in adult brain tissue, as well as with increased width of the left hemisphere central sulcus (Fig. 4B). Our comparative genomics analysis reveals that the trait-increasing allele (T) is a hominin-derived allele. We detect the same polymorphism at the site in the available ancient DNA data from Neandertals and Denisovans (Fig. 4C). Finally, we used The Human Genome Dating Atlas (HGD, Albers and McVean 2020) to estimate the age of the rs884370 polymorphism. HGD estimates that it is 54,171 generations old (95% confidence interval), corresponding to ∼1.35 mya assuming 25 years per generation.

## Discussion

We exploited summary statistics of a large-scale genome-wide scan for regional bilateral sulcal anatomy measures by Sun et al. (2022), which leveraged a dataset encompassing structural brain MRI and genotype information from over 26,530 individuals. We investigated the links between genetic variation underlying variability in sulcal organisation, and the marks that various evolutionary events have left in our genomes during human evolution. This non-invasive neuroimaging genetics strategy allowed us to reveal aspects of brain evolution by investigating the standing genomic and neuroanatomical variation in present-day humans, which has been largely unexplored by prior comparative primate neuroanatomical and neuronal model-based experimental work. Our study has three main advantages compared to prior similar studies: 1) larger sample size yielding higher power, 2) two previously unexplored sulcal measures (surface area and length), and 3) three enhanced evolutionary annotations of suitable SNP-coverage.

Our targeted approach tested 14 evolutionarily relevant sulcal measures and three annotations for SNP-heritability enrichment/depletion, but note that we could not predict whether to expect an enrichment or depletion signal in the Neandertal-ancestry related annotations, since the sulcal anatomy of Neandertals is largely unknown, due to absence of fossilized brain tissue. Our partitioned heritability analysis revealed significant enrichment signals in foetal brain HGEs for left and right hemispheric central sulcus width, which overlap with the findings of Lemaitre et al. (Lemaitre et al. 2023) showing right central sulcus heritability enrichment in foetal brain HGEs active in 7^th^ post-conception week. The central sulcus divides pre- and postcentral gyri, and the precentral gyrus, which is involved in motor-hand control in a contralateral manner (Hopkins et al. 2014). The morphological differentiation of this sulcus among primates does not show a lateralized pattern (Hopkins et al. 2014), which is in line with our robust enrichment findings for both hemispheres. We believe that the larger sample size of the current study (N=26,530), and the fact that our enhanced foetal brain HGE annotation covers a wider neurodevelopmental window of the foetal brain, as well as encompassing more SNPs, enabled us to identify a global (i.e. bilateral) enrichment signal for the central sulcus, as opposed to the right-hemispheric finding of Lemaitre et al. (2023). Significant enrichment signals for the left and right calloso-marginal posterior fissure in the aforementioned study are absent in our results, possibly because our foetal brain HGE annotation minimises the false-positive enrichment estimates. The significant heritability enrichment for the surface area of right parieto-occipital in foetal brain HGEs further supports our central sulcus finding, as this sulcus separates parietal and occipital cortex (Bruner 2018) regions hypothesized to show human specialisation for body-tool coordination (Hutchison et al. 2015; Sulpizio et al. 2016). Additionally, we identified significant heritability enrichment in foetal brain HGEs for the width of the superior temporal sulcus, which is known for its involvement in social perception and cognition, and deemed to have a key role in the convergence of spoken and written language (Deen et al. 2015; Wilson et al. 2018). Interestingly, this enrichment was detected in the right hemisphere, which might highlight the prominent rightward asymmetries in the central regions of the superior temporal sulcus, with particular differences in depth across hemispheres, which are less pronounced in chimpanzees (Leroy et al. 2015; Hopkins et al. 2023). Altogether, the enrichment signals in foetal brain HGEs for the entirety of the central sulcus, the right parieto-occipital sulcus, and the superior temporal sulcus may indicate the involvement of human-gained gene regulatory elements in the evolution of human hand dexterity, social cognition, and speech. Here, we note that in addition to the four sulci that we tested, there are other sulci that are relevant for human brain evolution. For example, paracingulate sulcus, the intralimbic sulcus, anterior and posterior vertical paracingulate sulcus, as well as dorso- and ventromedial-polar sulci were associated with the expansion of the medial frontal cortex in humans compared to chimpanzees (Amiez et al. 2021). However, these sulci were not part of the sulcus atlas used by Sun et al. (2022), and therefore could not be included in our targeted approach.

The exploratory analysis on the remaining 139 sulcal parameters revealed a robust significant heritability enrichment signal in the human-gained enhancers for the depth of the right olfactory sulcus. The olfactory sulcus divides the medial orbitofrontal gyrus and the rectus gyrus, harbouring the olfactory bulb and tract (Zang et al. 2020), suggesting a close link between the anatomies of the olfactory sulcus and the bulb. Reduced olfactory sulcus depth has been linked to congenital anosmia, which is a disorder identified by the inability of smell (Huart et al. 2011). On the other hand, it has been reported in the primate comparative neuroanatomy literature that a reduction in olfactory bulb size was followed by a shift towards visual system reliance, evolving independently in OWMs and apes (Gonzales et al. 2015). This is in line with our enrichment finding in foetal brain HGEs for the right olfactory sulcus, suggesting involvement of human-gained regulatory regions in the evolution of human olfactory structure. The fact that we detect this signal only for the right hemispheric olfactory sulcus is difficult to interpret. The right olfactory sulcus has been reported to be deeper compared to the left, showing an anatomic lateralisation for this sulcus. Intriguingly, olfactory regions have been shown to be functionally lateralised as well, with the left hemisphere being involved in emotional processing of odours, and the right side in the process of recognition memory (Royet and Plailly 2004). Combining our right hemisphere enrichment finding with the anatomical and functional lateralisation of the olfactory sulcus might suggest differential genetic roots for odour-based recognition memory in humans and OWMs.

Interpretation of the heritability depletion in archaic deserts for the width of left hemisphere internal frontal sulcus, inferior frontal sulcus, and posterior inferior temporal sulcus, as well as for the width of right hemisphere insula and the posterior terminal ascending branch of superior temporal sulcus is more speculative. Heritability depletion in archaic deserts might indicate that these genomic regions lack genes and regulatory elements involved in the aspects of foetal brain development which shape this particular sulcal structure. In line with our results, Alagöz et al. (2022) reported significant heritability depletion in archaic deserts for surface area of the left pars opercularis, which is topologically in close proximity to both the internal frontal sulcus and inferior frontal sulcus, and is associated with speech processing, articulation and phonological processing (Price 2012).

Finally, we focused on an example locus to show the intricate links between common genetic variants associated with sulcal morphology, functional context, and their evolutionary past. We chose a genome-wide significant locus (P<5×10^-8^) on chromosome 3 that is associated with the left hemisphere central sulcus width, tagged by the lead SNP rs884370. Our analysis showed that the rs884370 is located within a foetal brain HGE element near *ZIC4*, and regulates the neural expression of *ZIC1* and *ZIC4,* which are adjacent paralogue genes known to be involved in neurogenesis (Grinberg et al. 2004). We further demonstrate that the *ZIC4* promoter region interacts with this HGE element, providing evidence for the involvement of human-gained enhancers in the evolution of neurodevelopment through differential regulation of *ZIC4* across primates. A previous study (Alagöz et al. 2022) identified common genetic variants associated with the surface area of the left-hemisphere pars triangularis, a cortical region involved in speech and language, located in the same foetal brain HGE element that is an eQTL for *ZIC4* expression in human cortical tissue. The findings here converge with this previous finding, showing that the standing genetic variation on HGE elements impacts various aspects of human cortical anatomy from surface area to sulcal morphology. In the exploratory stream of the study, we did not identify significant heritability enrichments or depletions within Neandertal introgressed alleles for any sulcal traits. The absence of such signals may indicate that the common genetic variants associated with sulcal organization do not concentrate at or are diminished from the Neandertal introgressed regions. This is consistent with larger neuroanatomical differences across primates, rather than between *H. sapiens* and Neandertals. Our comparative genomic findings for rs884370 further support this by showing that this polymorphism is shared across present-day humans, Neandertals and Denisovans, but is not identified in non-human primates.

Our study has a number of limitations. First, the comparative neuroanatomy field has investigated sulcal structures at various levels, with higher priority on the sulci with clinical importance or co-localization with cortical regions such as Broca’s area. Thus, the breadth of prior information on sulcal anatomy is skewed towards a small subset of sulci. This makes it challenging to formulate testable hypotheses for the genetic and molecular evolution of sulcal organisation at a global level. Hence, we had to split our analytic pipeline into targeted and exploratory branches, each focusing on different sets of sulci with different levels of prior available knowledge. Second, this work is based on GWAS summary statistics (Sun et al. 2022) derived from a European ancestry population, hindering a full understanding of the link between human genetic variation and phenotypic variability in sulcal morphology. Third, even though we used the largest available GWAS of sulcal morphology with data from 26,530 people, we acknowledge that even larger sample sizes will be required to capture all true positive heritability enrichment and depletion signals in the tested annotations. Fourth, as with most large-scale neuroimaging studies, the quality of the MRI-based sulcal parameter estimates was not checked manually, due to practical challenges of doing this across tens of thousands of individuals. Fifth, in addition to three evolutionary annotations of the human genome that we tested here, there are other interesting sets of genomic regions that would be relevant for studying human brain evolution. However, as previously reported (Alagöz et al. 2022), annotations often cover a relatively small percentage of the SNPs available in population genetics reference panels, which can lead to spurious enrichment/depletion signals and significantly negative partitioned heritability estimates, beyond biological plausibility. We thus preferred to adopt a more conservative approach by using evolutionary annotations with at least 1% SNP-coverage, to minimise significant negative heritability estimates, and made no attempt to interpret such findings as they are not biologically meaningful. Despite our efforts to minimise such results, our analysis yielded a few significant negative heritability enrichment/depletion signals, which are not biologically interpretable. We believe that the discovery of more genomic regions of interest for primate brain evolution, and the availability of new sophisticated methods to study complex trait evolution will fuel the future of human brain evolution research.

In summary, here we shed light on the complex links between common genetic variants associated with various aspects of human sulcal organisation, and evolutionary annotations of the human genome such as foetal brain HGEs, Neandertal introgressed fragments, and archaic deserts. Our results demonstrate how integrating neuroimaging genetics with evolutionary annotations of the human genome provides a promising method for future human brain evolution studies by filling the gap between gross anatomical and *in vitro* neuronal model-based studies.

## Acknowledgements

O.G., G.A., B.M., and S.E.F. are supported by the Max Planck Society. O.G. is also supported by the German Federal Ministry of Education and Research (BMBF). The funders had no role in study design, data collection and analysis, the decision to publish, or the preparation of the manuscript. S.E.F. is a member of the Center for Academic Research and Training in Anthropogeny (CARTA). We thank Neda Jahanshad for providing the sulcal morphology GWAS summary statistics, and Else Eising for her critical feedback on functional annotation of the GWAS results.

## Data availability

GWAS summary statistics used in this study are publicly available at https://enigma-brain.org/sulcibrowser.

## Code availability

All scripts used for analyses are publicly available on the Gitlab repository: https://gitlab.gwdg.de/ole.goltermann/evolutionarysulcusmorph/-/tree/main/Analysis

## Author contributions

O.G., G.A., B.M., and S.E.F. designed research; O.G., G.A., and B.M. performed research; O.G. and G.A. analyzed data; G.A. wrote the initial draft of the manuscript; O.G., G.A., B.M., and S.E.F. provided critical feedback and commented on the manuscript. The authors declare no competing interest.

## References

Alagöz G, Molz B, Eising E, Schijven D, Francks C, Stein JL, Fisher SE. 2022. Using neuroimaging genomics to investigate the evolution of human brain structure. Proceedings of the National Academy of Sciences. 119(40):e2200638119. doi:10.1073/pnas.2200638119.

Albers PK, McVean G. 2020. Dating genomic variants and shared ancestry in population-scale sequencing data. PLOS Biology. 18(1):e3000586. doi:10.1371/journal.pbio.3000586.

Amiez C, Sallet J, Giacometti C, Verstraete C, Gandaux C, Morel-Latour V, Meguerditchian A, Hadj-Bouziane F, Ben Hamed S, Hopkins WD, et al. A revised perspective on the evolution of the lateral frontal cortex in primates. Sci Adv. 9(20):eadf9445. doi:10.1126/sciadv.adf9445.

Amiez C, Sallet J, Hopkins WD, Meguerditchian A, Hadj-Bouziane F, Ben Hamed S, Wilson CRE, Procyk E, Petrides M. 2019. Sulcal organization in the medial frontal cortex provides insights into primate brain evolution. Nat Commun. 10(1):3437. doi:10.1038/s41467-019-11347-x.

Amiez C, Sallet J, Novek J, Hadj-Bouziane F, Giacometti C, Andersson J, Hopkins WD, Petrides M. 2021. Chimpanzee histology and functional brain imaging show that the paracingulate sulcus is not human-specific. Commun Biol. 4(1):1–12. doi:10.1038/s42003-020-01571-3.

Amiez C, Wilson CRE, Procyk E. 2018. Variations of cingulate sulcal organization and link with cognitive performance. Sci Rep. 8(1):13988. doi:10.1038/s41598-018-32088-9.

Auton A, Abecasis GR, Altshuler DM, Durbin RM, Abecasis GR, Bentley DR, Chakravarti A, Clark AG, Donnelly P, Eichler EE, et al. 2015. A global reference for human genetic variation. Nature. 526(7571):68–74. doi:10.1038/nature15393.

Brouwer RM, Klein M, Grasby KL, Schnack HG, Jahanshad N, Teeuw J, Thomopoulos SI, Sprooten E, Franz CE, Gogtay N, et al. 2022. Genetic variants associated with longitudinal changes in brain structure across the lifespan. Nat Neurosci. 25(4):421–432. doi:10.1038/s41593-022-01042-4.

Browning SR, Browning BL. 2011. Population Structure Can Inflate SNP-Based Heritability Estimates. Am J Hum Genet. 89(1):191–193. doi:10.1016/j.ajhg.2011.05.025.

Bruner E. 2018. Human Paleoneurology and the Evolution of the Parietal Cortex. Brain Behav Evol. 91(3):136–147. doi:10.1159/000488889.

Bulik-Sullivan BK, Loh P-R, Finucane HK, Ripke S, Yang J, Patterson N, Daly MJ, Price AL, Neale BM. 2015. LD Score regression distinguishes confounding from polygenicity in genome-wide association studies. Nat Genet. 47(3):291–295. doi:10.1038/ng.3211.

Bycroft C, Freeman C, Petkova D, Band G, Elliott LT, Sharp K, Motyer A, Vukcevic D, Delaneau O, O’Connell J, et al. 2018. The UK Biobank resource with deep phenotyping and genomic data. Nature. 562(7726):203–209. doi:10.1038/s41586-018-0579-z.

Deen B, Koldewyn K, Kanwisher N, Saxe R. 2015. Functional Organization of Social Perception and Cognition in the Superior Temporal Sulcus. Cereb Cortex. 25(11):4596– 4609. doi:10.1093/cercor/bhv111.

Elliott LT, Sharp K, Alfaro-Almagro F, Shi S, Miller KL, Douaud G, Marchini J, Smith SM. 2018. Genome-wide association studies of brain imaging phenotypes in UK Biobank. Nature. 562(7726):210–216. doi:10.1038/s41586-018-0571-7.

Ernst J, Kellis M. 2012. ChromHMM: automating chromatin-state discovery and characterization. Nat Methods. 9(3):215–216. doi:10.1038/nmeth.1906.

Finucane HK, Bulik-Sullivan B, Gusev A, Trynka G, Reshef Y, Loh P-R, Anttila V, Xu H, Zang C, Farh K, et al. 2015. Partitioning heritability by functional annotation using genome-wide association summary statistics. Nat Genet. 47(11):1228–1235. doi:10.1038/ng.3404.

Giusti-Rodríguez P, Lu L, Yang Y, Crowley CA, Liu X, Juric I, Martin JS, Abnousi A, Allred SC, Ancalade N, et al. 2019. Using three-dimensional regulatory chromatin interactions from adult and fetal cortex to interpret genetic results for psychiatric disorders and cognitive traits. :406330. doi:10.1101/406330. [accessed 2022 Jun 13]. https://www.biorxiv.org/content/10.1101/406330v2.

Gonzales LA, Benefit BR, McCrossin ML, Spoor F. 2015. Cerebral complexity preceded enlarged brain size and reduced olfactory bulbs in Old World monkeys. Nat Commun. 6(1):7580. doi:10.1038/ncomms8580.

Grasby KL, Jahanshad N, Painter JN, Colodro-Conde L, Bralten J, Hibar DP, Lind PA, Pizzagalli F, Ching CRK, McMahon MAB, et al. 2020. The genetic architecture of the human cerebral cortex. Science. 367(6484):eaay6690. doi:10.1126/science.aay6690.

Grinberg I, Northrup H, Ardinger H, Prasad C, Dobyns WB, Millen KJ. 2004. Heterozygous deletion of the linked genes ZIC1 and ZIC4 is involved in Dandy-Walker malformation. Nat Genet. 36(10):1053–1055. doi:10.1038/ng1420.

Herculano-Houzel S. 2011. Not all brains are made the same: new views on brain scaling in evolution. Brain Behav Evol. 78(1):22–36. doi:10.1159/000327318.

Hopkins WD, Coulon O, Meguerditchian A, Staes N, Sherwood CC, Schapiro SJ, Mangin J-F, Bradley B. 2023. Genetic determinants of individual variation in the superior temporal sulcus of chimpanzees (Pan troglodytes). Cereb Cortex. 33(5):1925–1940. doi:10.1093/cercor/bhac183.

Hopkins WD, Meguerditchian A, Coulon O, Bogart S, Mangin J-F, Sherwood CC, Grabowski MW, Bennett AJ, Pierre PJ, Fears S, et al. 2014. Evolution of the Central Sulcus Morphology in Primates. BBE. 84(1):19–30. doi:10.1159/000362431.

Howe KL, Achuthan P, Allen James, Allen Jamie, Alvarez-Jarreta J, Amode MR, Armean IM, Azov AG, Bennett R, Bhai J, et al. 2021. Ensembl 2021. Nucleic Acids Research. 49(D1):D884–D891. doi:10.1093/nar/gkaa942.

Huart C, Meusel T, Gerber J, Duprez T, Rombaux P, Hummel T. 2011. The Depth of the Olfactory Sulcus Is an Indicator of Congenital Anosmia. AJNR Am J Neuroradiol. 32(10):1911–1914. doi:10.3174/ajnr.A2632.

Hutchison RM, Culham JC, Flanagan JR, Everling S, Gallivan JP. 2015. Functional subdivisions of medial parieto-occipital cortex in humans and nonhuman primates using resting-state fMRI. Neuroimage. 116:10–29. doi:10.1016/j.neuroimage.2015.04.068.

de Klein N, Tsai EA, Vochteloo M, Baird D, Huang Y, Chen C-Y, van Dam S, Oelen R, Deelen P, Bakker OB, et al. 2023. Brain expression quantitative trait locus and network analyses reveal downstream effects and putative drivers for brain-related diseases. Nat Genet. 55(3):377–388. doi:10.1038/s41588-023-01300-6.

Lemaitre H, Le Guen Y, Tilot AK, Stein JL, Philippe C, Mangin J-F, Fisher SE, Frouin V. 2023. Genetic variations within human gained enhancer elements affect human brain sulcal morphology. Neuroimage. 265:119773. doi:10.1016/j.neuroimage.2022.119773.

Leroy F, Cai Q, Bogart SL, Dubois J, Coulon O, Monzalvo K, Fischer C, Glasel H, Van der Haegen L, Bénézit A, et al. 2015. New human-specific brain landmark: The depth asymmetry of superior temporal sulcus. Proceedings of the National Academy of Sciences. 112(4):1208–1213. doi:10.1073/pnas.1412389112.

Lodewijk GA, Fernandes DP, Vretzakis I, Savage JE, Jacobs FMJ. 2020. Evolution of Human Brain Size-Associated NOTCH2NL Genes Proceeds toward Reduced Protein Levels. Mol Biol Evol. 37(9):2531–2548. doi:10.1093/molbev/msaa104.

Mafessoni F, Grote S, de Filippo C, Slon V, Kolobova KA, Viola B, Markin SV, Chintalapati M, Peyrégne S, Skov L, et al. 2020. A high-coverage Neandertal genome from Chagyrskaya Cave. Proceedings of the National Academy of Sciences. 117(26):15132–15136. doi:10.1073/pnas.2004944117.

Marzke MW. 1997. Precision grips, hand morphology, and tools. Am J Phys Anthropol. 102(1):91–110. doi:10.1002/(SICI)1096-8644(199701)102:1<91::AID-AJPA8>3.0.CO;2-G.

Mota B, Herculano-Houzel S. 2015. Cortical folding scales universally with surface area and thickness, not number of neurons. Science. 349(6243):74–77. doi:10.1126/science.aaa9101.

Patel GH, Sestieri C, Corbetta M. 2019. The evolution of the temporoparietal junction and posterior superior temporal sulcus. Cortex. 118:38–50. doi:10.1016/j.cortex.2019.01.026.

Price CJ. 2012. A review and synthesis of the first 20years of PET and fMRI studies of heard speech, spoken language and reading. NeuroImage. 62(2):816–847. doi:10.1016/j.neuroimage.2012.04.062.

Prüfer K, de Filippo C, Grote S, Mafessoni F, Korlević P, Hajdinjak M, Vernot B, Skov L, Hsieh P, Peyrégne S, et al. 2017. A high-coverage Neandertal genome from Vindija Cave in Croatia. Science. 358(6363):655–658. doi:10.1126/science.aao1887.

Prüfer K, Racimo F, Patterson N, Jay F, Sankararaman S, Sawyer S, Heinze A, Renaud G, Sudmant PH, de Filippo C, et al. 2014. The complete genome sequence of a Neanderthal from the Altai Mountains. Nature. 505(7481):43–49. doi:10.1038/nature12886.

Purcell S, Neale B, Todd-Brown K, Thomas L, Ferreira MAR, Bender D, Maller J, Sklar P, de Bakker PIW, Daly MJ, et al. 2007. PLINK: A Tool Set for Whole-Genome Association and Population-Based Linkage Analyses. Am J Hum Genet. 81(3):559–575.

Reilly SK, Yin J, Ayoub AE, Emera D, Leng J, Cotney J, Sarro R, Rakic P, Noonan JP. 2015. Evolutionary changes in promoter and enhancer activity during human corticogenesis. Science. 347(6226):1155–1159. doi:10.1126/science.1260943.

Ronan L, Fletcher PC. 2015. From genes to folds: a review of cortical gyrification theory. Brain Struct Funct. 220(5):2475–2483. doi:10.1007/s00429-014-0961-z.

Royet J-P, Plailly J. 2004. Lateralization of Olfactory Processes. Chemical Senses. 29(8):731–745. doi:10.1093/chemse/bjh067.

Schmitt AD, Hu M, Jung I, Xu Z, Qiu Y, Tan CL, Li Y, Lin S, Lin Y, Barr CL, et al. 2016. A Compendium of Chromatin Contact Maps Reveals Spatially Active Regions in the Human Genome. Cell Rep. 17(8):2042–2059. doi:10.1016/j.celrep.2016.10.061.

Schmitz MT, Sandoval K, Chen CP, Mostajo-Radji MA, Seeley WW, Nowakowski TJ, Ye CJ, Paredes MF, Pollen AA. 2022. The development and evolution of inhibitory neurons in primate cerebrum. Nature. 603(7903):871–877. doi:10.1038/s41586-022-04510-w.

Shadrin AA, Kaufmann T, van der Meer D, Palmer CE, Makowski C, Loughnan R, Jernigan TL, Seibert TM, Hagler DJ, Smeland OB, et al. 2021. Vertex-wise multivariate genome-wide association study identifies 780 unique genetic loci associated with cortical morphology. Neuroimage. 244:118603. doi:10.1016/j.neuroimage.2021.118603.

Sulpizio V, Committeri G, Lambrey S, Berthoz A, Galati G. 2016. Role of the human retrosplenial cortex/parieto-occipital sulcus in perspective priming. Neuroimage. 125:108–119. doi:10.1016/j.neuroimage.2015.10.040.

Sun BB, Loomis SJ, Pizzagalli F, Shatokhina N, Painter JN, Foley CN, Jensen ME, McLaren DG, Chintapalli SS, Zhu AH, et al. 2022. Genetic map of regional sulcal morphology in the human brain from UK biobank data. Nat Commun. 13(1):6071. doi:10.1038/s41467-022-33829-1.

Tashman KC, Cui R, O’Connor LJ, Neale BM, Finucane HK. 2021. Significance testing for small annotations in stratified LD-Score regression. :2021.03.13.21249938. doi:10.1101/2021.03.13.21249938. [accessed 2023 Oct 18]. https://www.medrxiv.org/content/10.1101/2021.03.13.21249938v1.

Tilot AK, Khramtsova EA, Liang D, Grasby KL, Jahanshad N, Painter J, Colodro-Conde L, Bralten J, Hibar DP, Lind PA, et al. 2021. The Evolutionary History of Common Genetic Variants Influencing Human Cortical Surface Area. Cerebral Cortex. 31(4):1873–1887. doi:10.1093/cercor/bhaa327.

Tocheri MW, Orr CM, Jacofsky MC, Marzke MW. 2008. The evolutionary history of the hominin hand since the last common ancestor of Pan and Homo. J Anat. 212(4):544–562. doi:10.1111/j.1469-7580.2008.00865.x.

Van Essen DC, Donahue CJ, Glasser MF. 2018. Development and Evolution of Cerebral and Cerebellar Cortex. Brain Behav Evol. 91(3):158–169. doi:10.1159/000489943.

Vernot B, Akey JM. 2014. Resurrecting surviving Neandertal lineages from modern human genomes. Science. 343(6174):1017–1021. doi:10.1126/science.1245938.

Vernot B, Tucci S, Kelso J, Schraiber JG, Wolf AB, Gittelman RM, Dannemann M, Grote S, McCoy RC, Norton H, et al. 2016. Excavating Neandertal and Denisovan DNA from the genomes of Melanesian individuals. Science. 352(6282):235–239. doi:10.1126/science.aad9416.

Watanabe K, Taskesen E, van Bochoven A, Posthuma D. 2017. Functional mapping and annotation of genetic associations with FUMA. Nat Commun. 8(1):1826. doi:10.1038/s41467-017-01261-5.

Willbrand EH, Voorhies WI, Yao JK, Weiner KS, Bunge SA. 2022. Presence or absence of a prefrontal sulcus is linked to reasoning performance during child development. Brain Struct Funct. 227(7):2543–2551. doi:10.1007/s00429-022-02539-1.

Wilson SM, Bautista A, McCarron A. 2018. Convergence of spoken and written language processing in the superior temporal sulcus. Neuroimage. 171:62–74. doi:10.1016/j.neuroimage.2017.12.068.

Zang Y, Whitcroft KL, Glöckler C, Hummel T. 2020. Is Handedness Associated with the Depth of the Olfactory Sulcus? ORL J Otorhinolaryngol Relat Spec. 82(3):115–120. doi:10.1159/000507787.

Zhao B, Zhang J, Ibrahim JG, Luo T, Santelli RC, Li Y, Li T, Shan Y, Zhu Z, Zhou F, et al. 2021. Large-scale GWAS reveals genetic architecture of brain white matter microstructure and genetic overlap with cognitive and mental health traits (n = 17,706). Mol Psychiatry. 26(8):3943–3955. doi:10.1038/s41380-019-0569-z.

Zheng J, Richardson TG, Millard LAC, Hemani G, Elsworth BL, Raistrick CA, Vilhjalmsson B, Neale BM, Haycock PC, Smith GD, et al. 2018. PhenoSpD: an integrated toolkit for phenotypic correlation estimation and multiple testing correction using GWAS summary statistics. Gigascience. 7(8):giy090. doi:10.1093/gigascience/giy090.

